# Predicted and Experimental NMR Chemical Shifts at Variable Temperatures: The Effect of Protein Conformational Dynamics

**DOI:** 10.1101/2023.01.25.525502

**Authors:** Xu Yi, Lichirui Zhang, Richard A. Friesner, Ann McDermott

## Abstract

NMR chemical shifts provide a sensitive probe of protein structure and dynamics. Prediction of shifts, and therefore interpretation of shifts, particularly for the frequently measured amidic ^15^N sites, remains a tall challenge. We demonstrate that protein ^15^N chemical shift prediction from QM/MM predictions can be improved if conformational variation is included via MD sampling, focusing on the antibiotic target, *E. coli* Dihydrofolate reductase (DHFR). Variations of up to 25 ppm in predicted ^15^N chemical shifts are observed over the trajectory. For solution shifts the average of fluctuations on the low picosecond timescale results in a superior prediction to a single optimal conformation. For low temperature solid state measurements, the histogram of predicted shifts for locally minimized snapshots with specific solvent arrangements sampled from the trajectory explains the heterogeneous linewidths; in other words, the conformations and associated solvent are ‘frozen out’ at low temperatures and result in inhomogeneously broadened NMR peaks. We identified conformational degrees of freedom that contribute to chemical shift variation. Backbone torsion angles show high amplitude fluctuations during the trajectory on the low picosecond timescale. For a number of residues, including I60, ψ varies by up to 60º within a conformational basin during the MD simulations, despite the fact that I60 (and other sites studied) are in a secondary structure element and remain well folded during the trajectory. Fluctuations in ψ appear to be compensated by other degrees of freedom in the protein, including φ of the succeeding residue, resulting in “rocking” of the amide plane with changes in hydrogen bonding interactions. Good agreement for both room temperature and low temperature NMR spectra provides strong support for the specific approach to conformational averaging of computed chemical shifts.

## 1. Introduction

NMR chemical shifts provide a sensitive probe of protein structure and dynamics ^1–3^. Unfortunately, prediction of shifts, and therefore interpretation of shifts, particularly for the frequently measured amidic ^15^N sites, remains a significant challenge. From quantum mechanics-based (QM) calculations, ^15^N chemical shifts are strongly influenced by local protein structure ^1,4,5^, including in particular, the backbone torsion angles ^6,7^ as well as electrostatic or hydrogen bonding effects.

A number of studies suggest that conformational averaging contributes to the complexity of shift interpretation ^8–10^. With hybrid quantum mechanics/molecular mechanics/molecular dynamics (MD-QM/MM), improved accuracy of calculations of ^15^N chemical shift tensors has been reported ^11^. Strong support for the importance of conformational averaging comes from cryogenic solid-state NMR studies of proteins for which significant NMR line broadening has been observed ^12–18^, consistent with freezing out disparate members of the ensemble. A number of studies support the hypothesis that broad linewidths in low temperature NMR are due to static conformational heterogeneity ^14–18^ including direct measurements of distribution in the local torsion angles^19^. This points to the possibility of characterizing the protein conformational ensemble using low temperature NMR linewidths.

In this study, we demonstrate the impact of conformational ensembles on NMR shifts, focusing on the antibiotic target, dihydrofolate reductase (DHFR), which has been well studied in terms of NMR chemical shifts and conformational dynamics. We focus on a specific site for which abundant experimental data are available, and which is evidently stably folded in a beta strand conformation, I60. Given that the observation of broad backbone amide ^15^N NMR linewidths at low temperature is relatively general we also make comparisons with other sites.

## 2. Results

### 2.1 QM/MM calculation based on MD simulation

For protein systems, hybrid quantum mechanics/molecular mechanics (QM/MM) has been applied with success previously. This approach results in increasingly reasonable computational demands, showing excellent and improving accuracy ^20^. Here the QM/MM approach was implemented with an automated fragmentation (AF) procedure, which was previously benchmarked with 20 proteins ^21^.

To investigate the importance of conformational sampling in NMR shifts, we computed an MD trajectory and selected samples from it to prepare an appropriate ensemble of native and near native conformations. This group of conformations was then studied using QM/MM to predict chemical shifts. A 1000 ns MD simulation was performed based on a solved DHFR:TMP X-ray crystallography structure ^22^. The backbone heavy atoms RMSD referenced to the initial state is shown in **Figure S7a**, demonstrating the overall stability of the whole protein during the MD trajectory. QM/MM calculations were used to study chemical shifts for snapshots extracted from the MD simulation. To study the effects of solvent, the locations and orientations of water molecules from the MD simulation were imported to AF-QM/MM. A DHFR:TMP X-ray crystallography structure ^22^ with a resolution of 2.5 Å and R-value of 0.23 was used for direct QM/MM calculation and MD simulation. Details of the MD simulation protocols and QM/MM calculation of the coordinates of each MD snapshot and X-ray crystal structure are presented in the Materials and Methods section.

To represent solution NMR data the predicted shifts were averaged over the ensemble. **Figure 1** shows the comparison between the chemical shift prediction from the X-ray crystallography structure ^22^ and that averaged over the conformations sampled with MD. Both the X-ray structure and the MD snapshots were energy minimized before QM/MM calculations. The correlation coefficient comparing experimental solution ^15^N shifts to single conformer (X-ray structure) predicted shifts is 0.69 and for Cα is 0.78. We then extracted 20 snapshots from the 1000 ns MD simulation, regularly sampled with an interval between snapshots of 50 ns. Before QM/MM calculation, energy minimization of each snapshot was performed, during which heavy atoms were constrained with harmonic restraints and hydrogen atoms were allowed to move. The minimization stopped when a full convergence was reached, or the heavy-atom RMSD was greater than 0.3 Å. After energy minimization, QM/MM calculations for each of the 20 snapshots using AF-QM/MM resulted in predicted average chemical shift with a better correlation to experimental solution state NMR shifts, 0.73 for N and 0.94 for Cα. By another metric, the RMSE (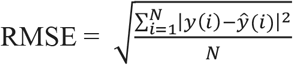, where N is the number of data points, y(i) is the i-th measurement, and *ŷ*(*i*) is its corresponding prediction.) can be thought of as a prediction accuracy, and was 4.8 ppm (^15^N) and 1.7 ppm (^13^Cα), respectively when averaging was included, vs 7.9 ppm and 3.6 ppm for the corresponding single conformation predictions. C’ QM/MM calculation results do not match the experimental chemical shift assignments as well as Cα (r = 0.51, RMSE = 2.3 ppm). Possibly the large CSA of the carbonyl, the resonance effects, and the important effects of hydrogen bonds are not well captured in the MD sampling. The improvement in the ^15^N shift prediction accuracy due to averaging is significant, especially given the importance of N shift measurements in protein studies.

**Figure 1:**
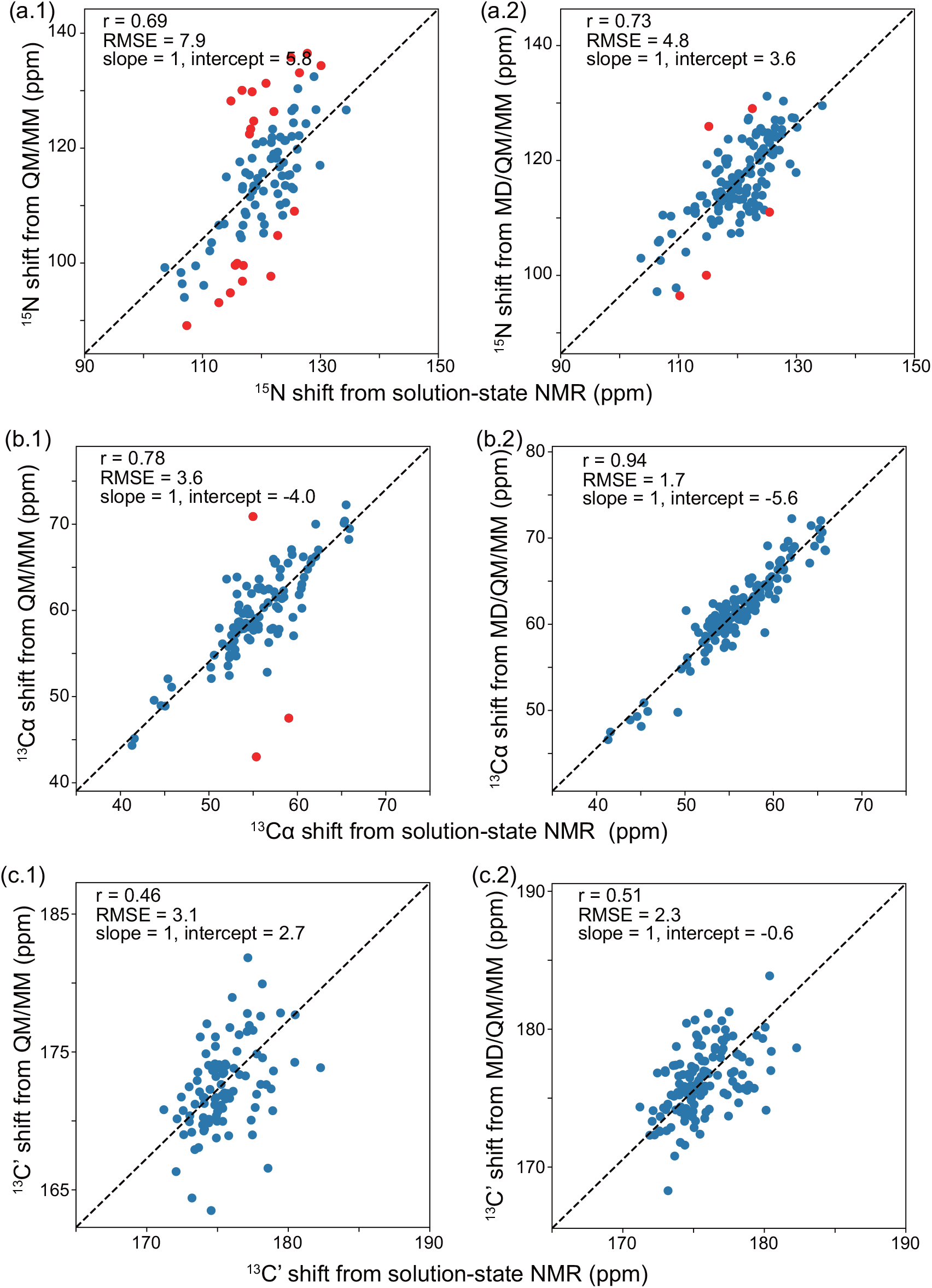
Correlation plots between experimental and QM/MM single X-ray diffraction conformer calculated shifts (a.1, b.1, c.1) or MD/QM/MM ensemble average calculated shifts (a.2, b.2, c.2) ^15^N (a), ^13^Cα (b), ^13^C’ (c) isotropic chemical shifts are shown. The experimental shifts were obtained with *E. coli* DHFR:TMP at 300 K on 800 MHz solution-state NMR spectrometer. The QM/MM calculation is based on an X-ray crystallography structure of *E. coli* DHFR:TMP ^22^. The MD/QM/MM calculation starts from a 1000 ns MD simulation of *E. coli* DHFR:TMP X-ray crystallography structure ^22^, followed by QM/MM calculations of 20 snapshots spaced by 50 ns intervals. All snapshots have water molecules preserved from the MD simulations, and restricted minimization was applied before QM/MM.

The shift prediction is relatively stable over the trajectory. Averaging snapshots from the different time ranges of the MD trajectory gives a similar chemical shift correlation with experimental results (**Figure S1**). The correlation is similar when extracting the snapshots from the first 200 ns, the last 200 ns, or the middle 200 ns in MD simulation. This result indicates that the whole MD calculation is stable, and the improvement due to hybrid QM/MM calculations and MD simulation is reproducible. Moreover, the key fluctuations that contribute to chemical shift averaging appear to operate on a relatively fast timescale consistent with the hypothesis that these fluctuations are averaged in the fast limit in solution NMR measurements.

In contrast with the narrow NMR spectra at room temperature, protein NMR peaks at low temperature are notably broad, putatively because various conformations freeze to create an inhomogeneous mixture. Since the QM/MM predicted shifts averaged over the MD simulation predict the solution-state NMR chemical shift at room temperature with good accuracy, we asked whether the ensemble of snapshots in the trajectory might be a good representation of the low temperature experimental ensemble. In order to address this question, we compared the amide ^15^N and ^13^C chemical shifts of each residue from MD snapshots with spectra collected at cryogenic temperatures. In **Figure 2**, the distribution of calculated ^15^N and ^13^C isotropic shifts in the snapshots matches well with the experimental contour at 100 K in solid-state NMR. Both the ^13^C chemical shift range and ^15^N shift range agree well, comparing the peak pattern and the QM/MM scatter points distribution. For the prediction of the five sites of IG-DHFR, 85% of the scatter points lay within the contours of DNP spectra. The chemical shifts were calculated with restricted minimization before QM/MM; without minimization, the predictions are less similar to the experiment, as shown in **Figure S2** where the ^15^N, ^13^C’ and ^13^Cα shifts uniformly distributed in the range of 95-140 ppm, 165-185 ppm and 53-68 ppm, respectively. The observation suggests that local minimization applied on MD snapshots before QM/MM allowing an adjustment with heavy atom RMSD < 0.3 Å is a reasonable simple model for experimental annealing / cooling of the sample.

**Figure 2:**
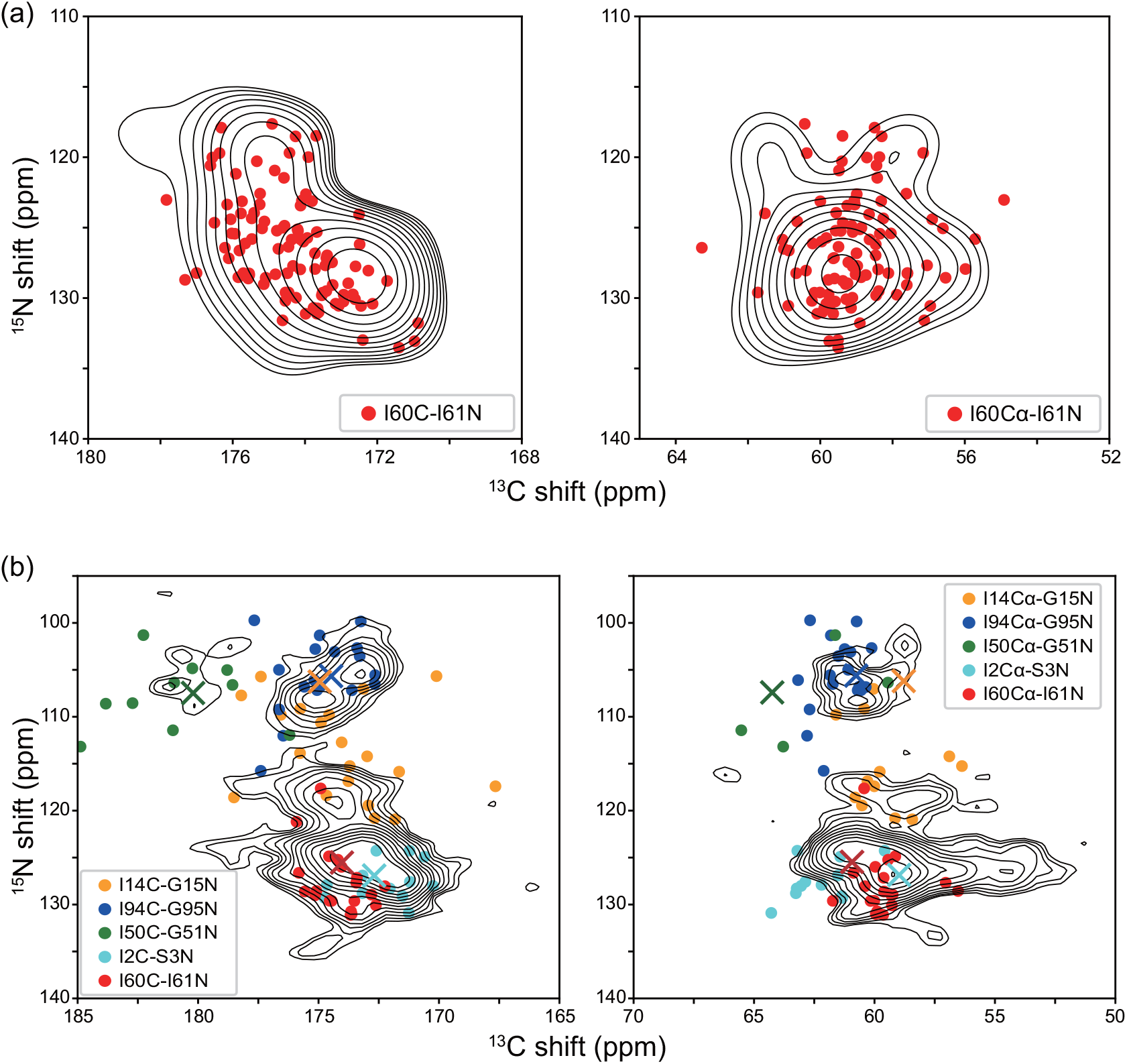
Low temperature solid-state NMR N-C correlation spectra of ^13^C,^15^N-Ile labeled *E. coli* DHFR:TMP (I-DHFR) (a) and ^13^C,^15^N-Ile, ^15^N-Gly labeled *E. coli* DHFR:TMP (IG-DHFR) (b) measured using DNP enhanced SSNMR at 105K. 160-I61 is the only residue pair expected in I-DHFR DNP spectra because of a sparse isotopic enrichment scheme (Ile only is enriched). Predicted chemical shifts for conformations sampled from MD and predicted with QM/MM are overlaid: 100 snapshots were minimized and calculated for I60-I61. In another enrichment scheme where both I and G are isotopically enriched “IG-DHFR” there are 5 residue pairs observed (Ile2-Ser3 (sheet), Ile14-Gly15 (coil), Ile50-Gly51 (helix), Ile60-Ile61 (sheet), Ile94-Gly95 (coil)). Their experimental lineshapes are compared with predictions based on 20 snapshots. Both C and N chemical shifts from QM/MM calculation were calibrated with the offsets from average chemical shifts of snapshots and solution-state NMR chemical shifts. The peak at Cα 53-55 ppm in IG-DHFR (b) results from a spinning sideband and was not predicted. The cross markers in (b) indicate the solution-state NMR chemical shifts for the 5 residue pairs.

### 2.2 Torsion angle variation in MD

The MD sampling studies described above suggest that NMR shifts fluctuate on a fast timescale at room temperature and are frozen into a broad inhomogeneous static distribution at cryogenic temperatures. The question arises as to the origins of these fluctuations in terms of conformational degrees of freedom. Inspecting the snapshots extracted every 10 ns from the MD trajectory, the torsion angles ψ and φ exhibit wide variations for many residues, as is shown in **Figure 3c and S3a**. We focus on Ile60-Ile61 which is located in the middle of a sheet structure in *E. coli* DHFR. The Ile60ψ value is 118° based on the X-ray crystallography structure of *E. coli* DHFR:TMP ^22^, and X-ray structures of *E. coli* DHFR complexes ^23^ in its catalytic cycle show variation for I60*ψ* in a range from 118° to 130°, indicating it as a relatively rigid site with small structural variation among different functional states. Although the ψ angle varies widely in the MD trajectory, Ile60 and Ile61 preserve beta-sheet structures throughout the trajectory, and the protein remains locally folded. The scatter points in **Figure 5** represent the φ-ψ correlation of these two residues in the Ramachandran plot. Such large fluctuations in backbone torison angles are consistent with stably folded structures if the change in one torsion angle is compensated by another, to keep the flanking reisues in the same position. Although ψ and φ both vary by 50 degrees in one amide, for example, Ile60-Ile61, the overall volume occupied by the protein chain is not much altered with these variations because of the correlation between ψ_i_ and φ_i+1_, which together result in the amide plane ‘rocking in place’ (Figure 3). Comparing the RMSD of atoms in residue Ile60 Cα vs. Ile60O (**Figure S7b-c**), Ile60Cα 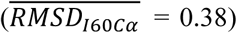 is more stationary, while the O atom makes somewhat wider excursions 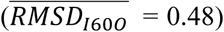. A strong correlation between ψ_i_ and φ_i+1_ is observed also for several other solvent inaccessible residues, **Figure S4 and Figure S11**, though some flexible solvent accessible residues showed more complex backbone dynamics. Similar observations in peptide ^24–26^ and protein ^27,28^ structures have been previously reported, where a dominant “crankshaft-like” motions in the plane of the amide results in a strong anti-correlated motion between φ_i+1_ and ψ_i_. It is noteworthy that other torsion angles such as the side chain torsion angle χ1_i_ also exhibit a variation up to 40° (**Figure S3b**), but do not correlate strongly with backbone torsion angles. The *ω*_i_ angle variation in the MD simulation within one amide is ∼ 7°, relatively small compared with other torsion angles, shown in **Figure S3c**.

**Figure 3:**
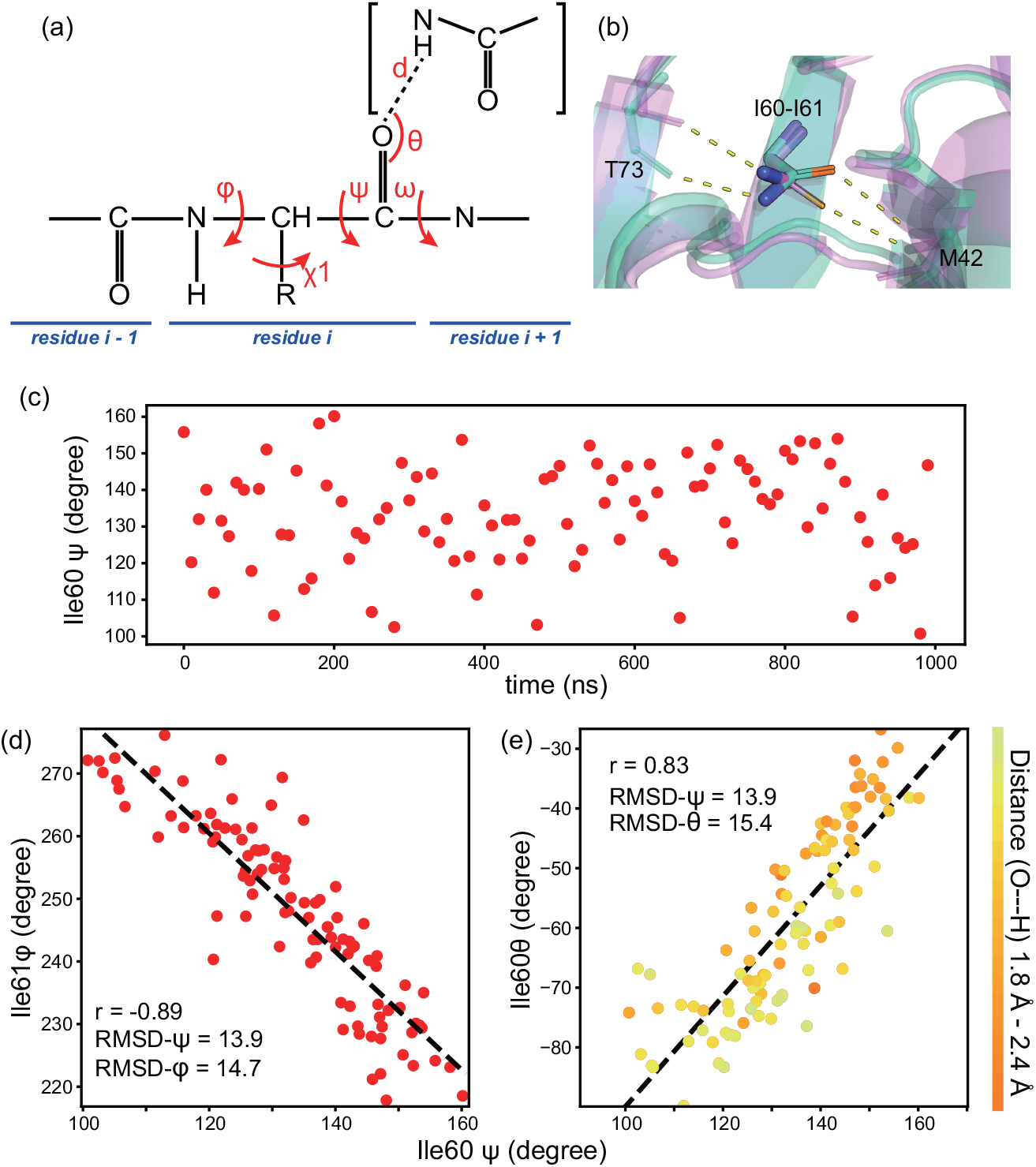
Protein structural variation during MD simulation at 300 K. (a) Dihedral angles and H bond angle/distance were analyzed. (b) Structure overlay of Ile60-Ile61 in snapshots at 200 ns in MD with Ile60ψ = 160° (green) and 160 ns in MD with Ile60ψ= 113° (pink) indicates that Cα position is unchanged while the N (blue) and carbonyl O (orange) move. The location of Ile60-Ile61 in the protein structure is shown in Figure 10. (c) The torsion angle ψ variation of Ile60 during the 1000 ns MD simulation is up to 60 degrees with snapshots extracted every 10 ns. (d) There is strong correlation between Ile61φ and Ile60ψ. Each scatter point represents 1 snapshot from MD simulation extracted every 10 ns. The correlation coefficient is -0.89. (e) The correlation plot of Ile60θ and Ile60ψ shows strong correlation between backbone torsion angle and H bond angle. H bond angle θ is defined as the angle between H bond (O---H) and C = O. Ile60 is located in a β -sheet structure and its O forms H bond with Met42 ^N^H in MD simulation. The closest water molecule is ∼7.5 Å away from the Ile60 O during the trajectory. The correlation between ψ_i+1_, φ_i_ and H bond angle θ_i_ is not unique for I60-I61, as shown in **Figure S11**.

### 2.3 Correlation between N isotropic chemical shifts and torsion angles

Numerous physical effects or structural features such as torsion angles are expected to influence ^15^N chemical shifts. We specifically focused on the implications of the fluctuation of ψ_i_ on the N_i+1_ chemical shift for Ile60-Ile61 in *E. coli* DHFR (**Figure 4a)**. This was partly motivated by the broad range of values for ψ observed in the MD simulations. Also, previous low temperature DNP-enhanced NMR experimental data show a correlation between this torsion angle and ^15^N shifts (Figure 4a of ^19^ ) A strong correlation between the I61 ^15^N chemical shifts and I60ψ values is observed from 100 snapshots in the 1000 ns MD trajectory. More importantly, both the isotropic chemical shifts and the torsion angles from MD/QM/MM agree with the experimental data. These data support the suggestion that variations in backbone torsion angles within the beta basin can have strong effects on the ^15^N chemical shifts.

**Figure 4:**
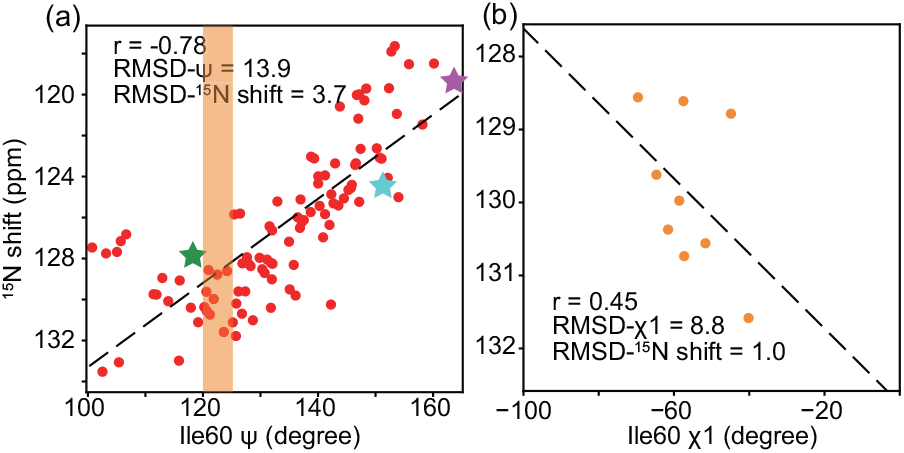
(a) QM/MM calculated Ile61N isotropic shifts are plotted as a function of I60ψ from 100 minimized snapshots from MD trajectory (red, circle). Three ψ values were measured experimentally at three ^15^N shift positions for Ile60 in low temperature DNP-enhanced NMR experiments^19^ and are shown here depicted with stars. The correlation between ψ angles and chemical shifts from both prior experimental and these computational results match well. The sidechain torsion angle I60χ1 were plotted with ^15^N shifts for snapshots with similar backbone torsion angle ψ, 120°< I60ψ <125° (b). Ile61N shifts vary up to 5 ppm with a 5° range of I60ψ, and I60χ1 is likely to contribute to this variation.

**Figure 5:**
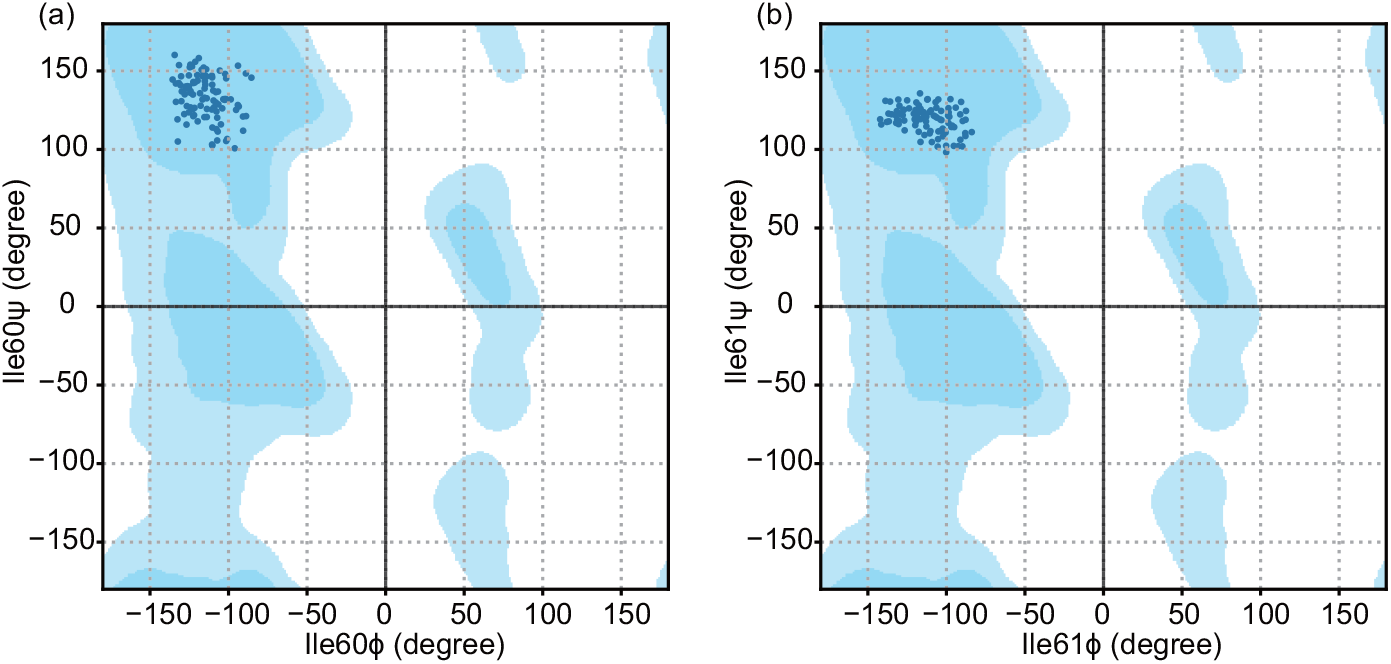
The φ-ψ of Ile60 (a) and Ile61(b) from snapshots extracted every 10 ns from 1000 ns MD simulation are shown with scatter points in the Ramachandran plot. Both residues exhibit torsion angles φ and ψ typical for a sheet structure, consistent with the X-ray crystallography structure of DHFR:TMP ^22^, though the range of values for 60 is notably broader.

The observation of wide variation in both the torsion angle and the ^15^N shift is not unique for Ile60-Ile61; as shown in **Figure S5**, most residues also exhibit a broad torsion angle variation of 40° or greater. Most residues also show predicted ^15^N shift variation over 10 ppm, consistent with the broad lines observed for all residues in low temperature NMR. The correlation of ψ with ^15^N chemical shifts is apparent for many other residues, as shown in **Figure S5d-f**, although it is not evident for every residue (**Figure S5a-c**). It is likely that the correlation is apparent for Ile60-61 because of the particularly broad range in ψ values sampled by Ile60. The effect of ψ on Ile61 ^15^N shift is most evident for conformers where the backbone makes excursions out of the most populated sheet basin around ψ =120*°* to conformations with ψ >130 *°*. Typical sheet /strand residues with ψ of 100°-130° show ^15^N shifts of 127 +/-6 ppm. By contrast, sheets that are closer to flat extended conformations with ψ > 150° exhibit shifts of 115 +/-5 ppm (**Figure 4a)**. Beta strand residues with more restricted ranges of torsion angle ψ from 90*°*-130*°* degrees, such Y111, do not show a significant correlation between backbone torsion ψ and ^15^N shift, and show a narrower span of both ψ and ^15^N shift (**Figure S5a**). A master plot QM/MM calculated ^15^N shifts of all the residues from 100 snapshots with their associated torsion angle ψ (**Figure S8)** shows that for beta sheets the effect of ψ on ^15^N chemical shifts is most pronounced for ψ ∼130-170°, while for sheets with ∼90-130°, the dependence is shallow and not monotonic. Ile60-Ile61 is located close to the cofactor binding site, which is empty in the structure that we studied. Possibly the particularly high flexibility of Ile60 is because of this “cavity”.

Analogously, the well populated basin for helices (ψ range -50°∼ 0°) has a mean ^15^N shift of 116 +/-5 ppm. Helical residues including 3, 109 and 149 have a relatively flat value for the ^15^N shift as a function of ψ in the range 0 to 60*°* with poorer correlation and a narrower span of both ψ and ^15^N shift.

Deviation in ^15^N chemical shifts within a defined range of ψ are as large as 10 ppm, indicating that other structural degrees of freedom besides ψ also contribute to variation in shift. The sidechain torsion χ1 has been discussed in relation to influencing backbone chemical shifts and therefore was also examined for correlations with ^15^N and ^13^C ^1,29,30^. We identified conformations from within the MD trajectory with ψ (120° < I60ψ < 125° in **Figure 4b**, 150°< I60ψ <155° in **Figure S9b**, 135° < I60ψ < 140° in **Figure S9c**). Within each group ^15^N chemical shifts vary by more than 3 ppm and sidechain torsion angles I60χ1 varied by up to 30° while staying within the typically preferred ‘gauche’ basin. The correlation plot suggests that χ1 is another factor influencing ^15^N shifts. The situation is more complex for solvent exposed and flexible residues, such as His149 (**Figure S10**), that show very broad ranges in other torsion angles (including ψ_i-1_ and ϕ_i_) that are not as well compensated by or correlated to other backbone angles. Consequently, they exhibit broad ranges of ^15^N_i_ shifts that are not evidently correlated to any one torsion angle (with a span of 25 ppm).

We explored whether the effect of ψ on the ^15^N shifts might occur because of changes in hydrogen bonding. Variation of backbone torsion angles and rocking of the amide plane described above is expected to modulate the interactions with hydrogen bond partners. This variation is of interest and is expected to have implications for energetics particularly dielectric and electrostatic characteristics. The hydrogen bond between the backbone NH and neighboring groups, as well as the hydrogen bond between the CO and other neighboring groups is also expected to N and C’ chemical shifts; both N and C chemical shift tensors are sensitive to the hydrogen bond variations (**Figure S6d-f**), especially for C’ atoms, consistent with previous literature ^31–33^. We investigated the hydrogen bond distance and angle over typical snapshots from the 1000 ns MD trajectory. For desolvated residues, such as Ile60-Ile61 in the middle of a sheet structure (**Figure 3b**), the identity of the protein residues that serve as hydrogen bond partners remain constant, and the pairwise trajectory was analyzed in terms of the distance and angle of the hydrogen bond. The shortest (or only) hydrogen bond partner for Ile60O is Met42 ^N^H. This distance varies by 0.6 Å over the trajectory. The shortest (or only) H bond partner of Ile61NH is the carbonyl O of Thr73. The correlation between the Ile61N chemical shifts and Ile60O---H distance, shown in **Figure S6a-b**, is not strong, though the hydrogen bond angle *θ* (relative to the CO vector of I60) shows a strong correlation with the torsion angle ψ. Analysis of solvent-accessible residues is complex because more than one water molecule is normally located within 4 Å of the amide atoms. Overall it is likely that the strong variation in hydrogen bonding environment is a major reason that ψ influences the ^15^N shift.

### 2.4 Timescale analysis of backbone torsion angle dynamics

In order to assess the timescale of backbone fluctuations, we analyzed a 100 ps segment of the MD simulation with snapshots extracted every 100 fs. The variation of torsion angles for Ile60ψ is shown in **Figure 6a**. The angle ranges from 90° to 150°, which is similar to the range observed in the longer time MD simulation shown in **Figure 3d**. Two states were defined based on the torsion angle range, State 1 with Ile60ψ > 125° and State 0 with Ile60ψ < 125°. The structure of Ile60-Ile61 amide undergoes a dynamic and structural change at ∼50 ps; before this time State 0 (**Figure 6b**) was preferable with Ile60ψ smaller than 125°, and after that time, State 1 (**Figure 6c**) with larger ψ was much more favored. Thus, Ile60ψ is considered as switching between two states, shown in **Figure 7a**. The timescale of this exchange motion is then determined from the time when Ile60ψ stays in either state continuously. In the first 50 ps, State 0 occupied more time with a residence time of 0.89±0.18 ps (**Figure 7d**), while the residence time of State 1 is 0.34±0.05 ps (**Figure 7b**). In the second half 50 ps, the residence time is 0.23±0.02 ps (**Figure 7e**) for State 0. The State 1 is more preferable and its residence time is longer than that can be extracted from this MD simulation. This dynamic timescale agrees with previous research about polypeptides ^24^, in which the anticorrelated motion of ψ_i_ and φ_i+1_ is reported to be on the timescale of 0.1 ps. This timescale was associated with protein backbone motions and intermolecular vibrations of the water hydrogen bond network ^34^, and these fast timescale atomic fluctuations were observed to facilitate large-scale, slower motions during enzyme catalysis ^35^.

**Figure 6:**
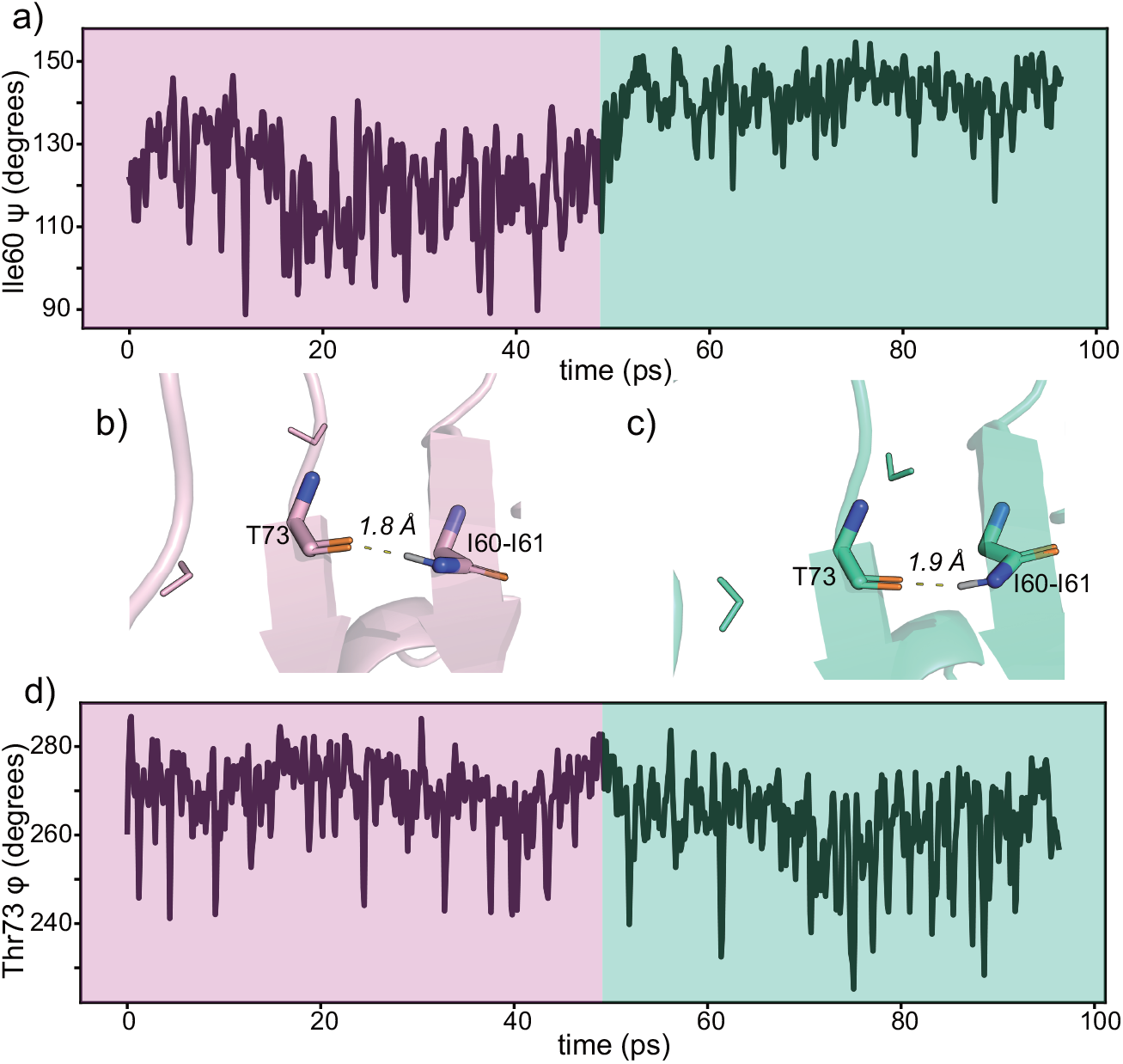
(a) The torsion angle ψ variation of Ile60 during the 100 ps MD simulation is ∼ 60 degrees, with snapshots extracted every 100 fs. Two main conformations were observed in the Ile60-Ile61 amide during this trajectory. Two structures were captured at 24 ps (b, pink) and 76 ps (c, green) as representatives of the two conformations with Ile60ψ of 108° and 146°, respectively. The pink structure is more dominant in the first 50 ps, while the green one is more favorable after 50 ps. The hydrogen bonds between the carbonyl O of Thr73 and Ile61NH were shown in yellow; N atoms are colored blue and backbone O atoms are colored orange. Thr73 was partially solvated with two water molecules (shown in sticks) within 3.5 Å from either Thr73N or Trp74N. (d) A Similar torsion angle variation was observed for Thr73φ.

**Figure 7:**
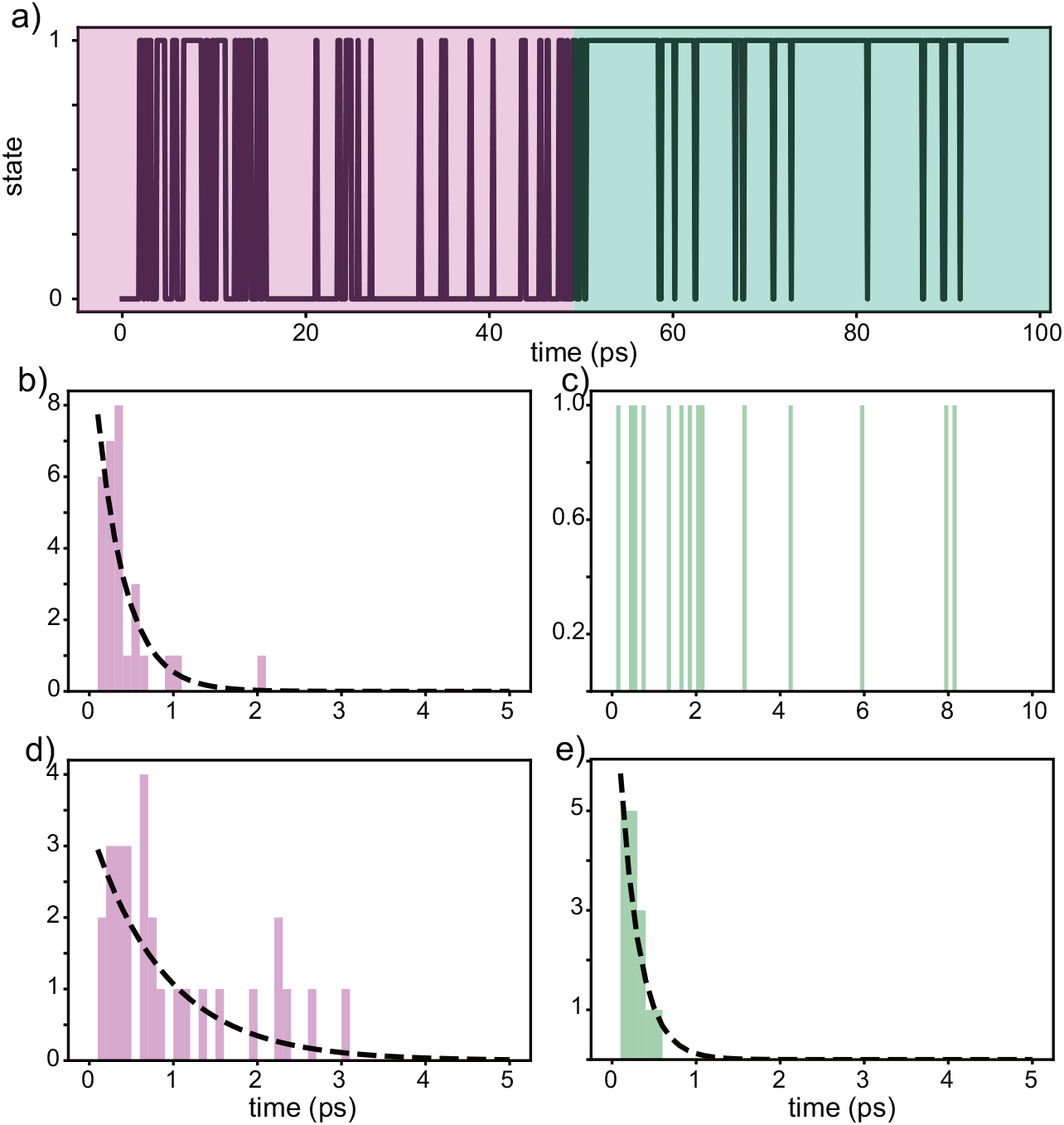
(a) Two states were defined based on Ile60ψ variation, State 1 (Figure 6c) with Ile60ψ > 125° and State 0 (Figure 6b) with Ile60ψ < 125°. State 0 is more dominant before (pink) 50 ps and State 1 after (green) 50 ps. The histograms of the residence times for Ile60ψ staying in State 0 (d, e) or State 1 (b, c) before (b, d, pink) and after (c, e green) 50ps were fit with a single exponential function (black dash line). For the first 50 ps trajectory, the residence time is 0.89±0.18 ps for State 0 and 0.34±0.05 ps for State 1. After 50 ps, the residence time is 0.23±0.02 ps for State 0. The residence time for State 1 is longer than that can be identified through this trajecotry.

The conformational variation of Ile60-Ile61 is likely to be influenced by its partner Thr73, whose O forms a hydrogen bond with Ile61^N^H. Thr73 located in another *β*-sheet structure and partially solvated, flips conformations at 50 ps, and probably causes the change in its partner Ile60. Thr73 in turn might be affected by the solvent. It forms closest hydrogen bonds with protein residues (Thr73O---Ile61^N^H, Thr73^N^H---Asn59O), yet there are also typically water molecules within 3.5 Å of this residue (**Figure 6b** and **6c**). Because the *β*-sheet is solvent-exposed, neighboring Trp74O and Val72O both form the shortest hydrogen bonds with water molecules. It is likely that the solvent interactions lead to increased structural variation for Thr73, which then cause it to influence nearby desolvated residues such at Ile60-61 through hydrogen bonding interactions. Longer MD simulations (**Figures 3c** and **S 14)** indicate that the two-state flipping happens regularly and reversibly throughout the trajectory.

## 3. Discussion

In summary, for relatively buried and structured residues in globular proteins, picosecond timescale large angle fluctuations in backbone torsion angles leads to large dynamic variation in backbone shifts. The changes, despite being over 60 degrees in amplitude, can be consistent with stably folded structures because they are compensated by other torsion angles, and the structure remains in the beta basin. The fluctuations should be considered when predicting room temperature shifts. They also contribute significantly to the inhomogeneous low temperature linewidths, because the conformations appear to be static (or interconvert very slowly) at 100K.

The large variation of torsion angles we observed in the MD trajectory may be surprising. Based on our analysis of MD snapshots, it is likely that the compensations from other backbone torsion angles and hydrogen bond angles preserve the global protein secondary structures. Analogous large amplitude backbone motions have been discussed previously in peptides ^24,36^, polymers ^37^, and proteins ^38,39^ with MD simulation in explicit or implicit solvent. In recent research of protein dynamics ^40^, several high-resolution protein structures were chosen for analyzing the amplitudes of protein motions; the (φ, ψ) distribution of Lys83 captured during dynamics of mouse protein kinase A is up to 150°, and the drastic changes are enabled by minor adjustments to the backbone geometry including bond lengths and angles. Similarly, the conformational transitions observed in Ala_7_ peptide are accomplished by a large change (64.4°) in the φ angle of the fourth residue; however, the entire chains are nearly perfectly superimposed before and after the torsion angle variation except for the atoms involved in ψ_4_, φ_4_ or ψ_3_ ^24^. The observation of thermally elicited large angle local torsion angle variations on the pico-second timescale is expected to have many experimental and computational implications. If the range of backbone torsion angles is as broad as is indicated here, it is likely that it would have implications for protein modeling, for solution NMR protein dynamics studies, and for fitting electron density in X-ray diffraction and other structure methods ^41^. Variation and interdependence of the backbone dihedral angles ^25–28^, was predicted to be associated with isotropic chemical shifts ^7,42^, NMR cross-relaxation ^43,44^ and scalar coupling constants ^45^.

We asked how fast, local large amplitude conformational fluctuations affect the chemical shifts of proteins, focusing on room temperature experimental solution shifts which reflect the average over the fluctuations. Although the empirical methods, like SHIFTX2 ^46^ and SPARTA+ ^47^, can predict chemical shifts with high accuracy, leading to correlation coefficients of 0.98 (^15^N), 0.99 (^13^C*α*), and 0.96 (^13^C’) with RMSE of 1.12, 0.44 and 0.53 ppm compared to the experimental results in 61 proteins ^46^, their usage can be limited to frequently observed and thermally averaged conformations, and so might be ill suited to sampling thermal excursions during dynamics ^48^. Fragment density functional theory calculation of NMR chemical shifts for protein, on the one hand, has become increasingly practical in terms of computational time ^21^ and, on the other hand, can capture nuanced variations in the structure including rare events and provide accurate and physics-based predictions. The chemical prediction of GB3 using the AF-QM/MM resulted in mean unsigned errors (MUE) of 4.8 ppm with R^2^ of 0.71 for N and 2.1 ppm with R^2^ of 0.79 for C*α* ^49^. Polenova’s group compared the QM/MM calculated chemical shifts of galection-3C with its MAS NMR chemical shifts ^50^. The correlation coefficient comparing predicted vs. observed shifts for ^13^C*α* and ^15^N was 0.83 and 0.68 using an X-ray structure determined at 100 K; a structure determined at 293 K gives a similar correlation, which is 0.84 for ^13^C*α* and 0.65 for ^15^N. Moreover, the explicit solvent molecules in QM/MM allow the interaction between the solvent and the protein to be calculated ^51–53^. An explicit–implicit solvation model with only one explicit water molecule per amine and amide proton is sufficient to yield accurate results for ^1^H (MAD ∼ 0.3 ppm) and for ^13^C (MAD ∼3 ppm) ^54^. Here, we conclude that it is possible to simulate the distribution in chemical shifts at room temperature and at low temperatures using QM/MM calculations, assuming that the following technical details and procedures are followed: 1) Averaging the QM/MM calculated shifts over a long enough MD trajectory to obtain better prediction at room temperature. 2) Applying restricted minimization on room temperature MD snapshots before QM/MM to mimic the freezing out protein conformation distribution. Conformational averaging has been observed to be necessary for QM/MM prediction ^51,52,55^. Previously, QM/MM prediction of a 32-amino-acid-long polypeptide averaging 500 snapshots from a 5 ns explicit solvent MD simulation gives MUE of 14.6 ppm with R^2^ value of 0.81 for ^15^N and MUE of 5.2 ppm with R^2^ value of 0.99 for ^13^C (not only ^13^C*α*). With our procedure, we obtained a correlation for ^13^C*α* (r = 0.94) and ^15^N (r = 0.73) atoms and RMSE of 4.8 ppm, 1.7 ppm, and 2.3 ppm for ^15^N, ^13^C*α*, and ^13^C’, respectively, which is significantly improved relative to the single conformation case where ^13^C*α* (r = 0.78) and ^15^N (r = 0.69), comparable to previous reports. Thus, we conclude that, while more work remains to be done, conformational averaging improves predictions of solution shifts significantly.

We asked how the fluctuations in the shift might affect low temperature NMR measurements. Combining the results of QM/MM calculated shifts with the snapshots from the MD simulation, we noticed that the ^15^N chemical shifts of protein exhibit a broad range. The variation can have a number of structural origins. The contribution of the backbone ^7,56,57^ and side-chain ^2,29,58^ torsion angles to ^15^N isotropic chemical shifts has been previously discussed; thus, their effects were carefully considered for a more accurate chemical shift prediction. Previous work indicates that high solvent accessibility correlates with high line widths in low temperature DNP-enhanced NMR spectra. As indicated in the previous literature ^59^, and from the QM/MM predictions reported here, the contribution of the hydrogen bonds to both ^15^N and ^13^C chemical shift tensors is large and the effect of isotropic shifts can be up to 8 ppm to ^15^N chemical shifts in model systems, and smaller for ^13^C isotropic shifts (< 2 ppm for ^13^C’ and < 0.5 for ^13^C*α*). For ^15^N chemical shift tensors with the hydrogen bond, δ_11_ appears to be sensitive to the presence of hydrogen bonding interactions due to its unique orientation in the molecular frame ^31^. For ^13^C chemical shift, the tensor components depend strongly on the hydrogen-bond length and angle, particularly δ_22_ ^60. 12^. These effects may explain the empirical trends regarding solvent accessibility, and including solvent interactions in chemical shift calculation, especially anisotropy prediction, is expected to be important ^29,33,61^. Accordingly, we chose to retain water molecule locations from MD simulations and to apply restricted minimization of the MD snapshots, to simulate the effects of sample freezing and predict inhomogeneous conformational distributions and inhomogeneous chemical shift distributions in the frozen protein.

The research reported here supports the hypothesis that much of the variation in chemical shifts and broad NMR linewidths for frozen samples is due to local conformational variation, where various backbone conformers are frozen into an inhomogeneous distribution at 100K (with interconversions on a timescale slower than milliseconds) ^14–18^. By contrast, they are averaged into a homogeneous distribution at room temperature (with interconversions on the sub-nanosecond timescale). This conclusion agrees well with low temperature experimental measurements of distributions in backbone ψ and correlations to the chemical shifts in various members for the ensemble. Backbone torsion angles, specifically correlated changes in ψ_i_ and φ_i+1_ and a resulting motion of the amide plane, are associated with these changes in NMR chemical shifts. While other structural degrees of freedom are likely to be variable and important as well, we showed here that for the N isotropic shift, ψ is a significant cause of variation.

## 4. Conclusions

We explored the connection between dynamics predicted by MD and NMR observables and presented strong indications of elevated thermal motions of the backbone of proteins. For liquid samples, averaging the QM/MM shift prediction over the trajectory improves the chemical shift prediction accuracy compared to predictions based on single global minimum conformations. The distributions in predicted values also agree well with the breadth of low temperature NMR spectra. Variation in the backbone torsion angles psi specifically appears to have marked effects on the ^15^N isotropic shifts.

## Materials and Methods

### Molecular dynamics simulations

Starting from the DHFR:TMP x-ray crystallography structure, the system was prepared using Protein Preparation Wizard in Schrödinger’s Maestro ^62^. Specifically, bond orders were assigned, and hydrogen atoms were added. The ionization state of TMP and ions suitable for pH 7.0 ± 1.0 were predicted using Epik ^63,64^. The H-bond network was then optimized with protonation states of residues at pH 7.5, as determined via pKa prediction by PROPKA ^65^. Finally, the structure was subjected to restrained energy minimization using the OPLS4 force field ^66^, with a heavy atom RMSD of 1.0 Å.

For MD simulations, the system was set up using System Builder in Maestro. The structure was placed in an orthorhombic box of TIP3P (36) water molecules with the box boundary at least 10 Å away from any protein atoms. 19 Na^+^ ions were added to neutralize the system. The total number of atoms in the MD system is 20,141. The MD simulations were carried out using Desmond ^67^ with the OPLS4 force field. Simulations were kept at a constant temperature of 300 K and a constant pressure of 1.01325 bar maintained with the Nosé-Hoover chain thermostat (*τ*=1 ps)^68^ and the Martyna-Tobias-Klein barostat (*τ*=2 ps) ^69^, respectively. The integration time-step was set to 2 fs, 2 fs, and 6 fs for bonded, near, and far interactions, respectively. The particle-mesh Ewald (PME) method was used for electrostatic interactions, and the cutoff value for the short-range contribution was set to 9 Å. The system was relaxed before the simulation, using the default relaxation protocol in Desmond. The total simulation time was 1 microsecond, and snapshots were generated after every 1 nanosecond, resulting in 1000 snapshots in total.

### AF-QM/MM calculations

The isotropic chemical shifts of backbone ^15^N and ^13^C were calculated using the AF-QM/MM method with the AFNMR package ^21^. In AFNMR, the protein is divided into individual residues. Each residue and its surrounding atoms (< 3.4 Å) are treated with density functional theory (DFT), and the remaining atoms are described by atomic point charges computed by a Poisson-Boltzmann procedure with the Amber force field using the “solinprot” program from the MEAD package. The dielectric constants were set to be ε = 1 for the quantum region, ε = 4 for the remainder of the protein, and ε = 78 for the solvent region in the Poisson-Boltzmann equation. The Amber force field parameters for the ligand were obtained using the Antechamber package in AmberTools.^70^ The snapshots were first refined with restrained energy minimization using the OPLS4 force field, with a heavy atom RMSD cutoff of 0.3 Å. In restrained minimization, the structure was subjected to conjugate-gradient minimization with harmonic restraints on heavy atoms. Minimization stops when a full convergence is reached, or the heavy-atom RMSD exceeds the cutoff. Then the QM/MM calculation input file for each residue was generated by AFNMR. QM/MM calculations were performed using ORCA ^71^ with the OLYP functional and the psSseg-0 basis set ^72^ optimized for chemical shift calculations. Calculated chemical shifts were referenced using the standard shielding constants (tetramethylsilane for ^13^C and nitromethane for ^15^N) calculated at the same level of theory.

The initial structures and the configuration files describing additional technical details of MD simulations and QM/MM calculations are available at https://github.com/zlcrrrr/DHFR_MD_QMMM. <u>The MD trajectory, minimized snapshots, and predicted chemical shifts are available at</u> https://osf.io/wqyb4/.

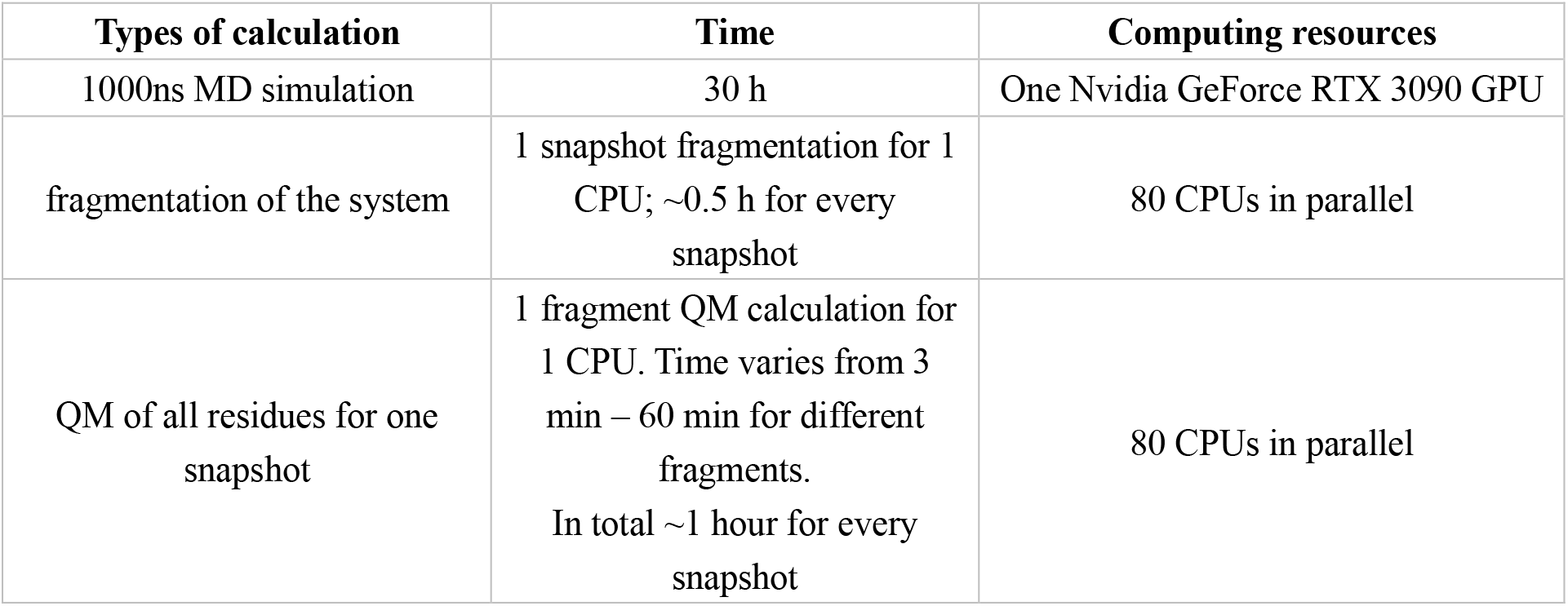

| **Types of calculation** | **Time** | **Computing resources** |
| --- | --- | --- |
| 1000ns MD simulation | 30 h | One Nvidia GeForce RTX 3090 GPU |
| fragmentation of the system | 1 snapshot fragmentation for 1 CPU; ~0.5 h for every snapshot | 80 CPUs in parallel |
| QM of all residues for one snapshot | 1 fragment QM calculation for 1 CPU. Time varies from 3 min – 60 min for different fragments.<br>In total ~1 hour for every snapshot | 80 CPUs in parallel |

## Acknowledgements

This work was supported by a grant from the NSF (MCB 1913885), and a Biomedical Technology Development and Dissemination Center grant from the NIH (1RM1GM145397-01). We thank David Case and Yunyao Xu for helpful discussions.

