## Supplementary Figures for "Predicted and Experimental NMR Chemical Shifts at Variable Temperatures: The Effect of Protein Conformational Dynamics"

**Supporting information for**

### Protein structural heterogeneity of fast timescale dynamics captured by low temperature NMR

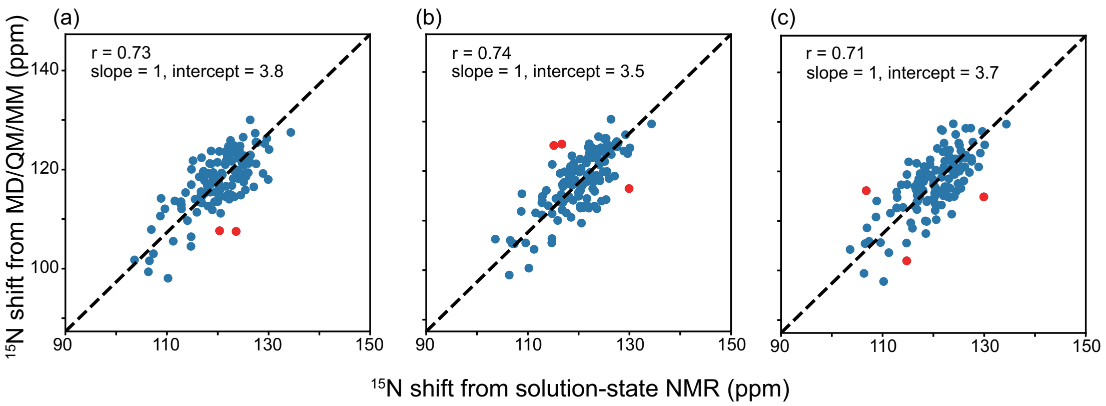


**Figure S1:** The correlations between experimental and MD/QM/MM calculated $^{15}$N chemical shifts are consistent in different time range of one MD trajectory. 20 snapshots were extracted from (a) 1-3 ns, (b) 4-6 ns, and (c) 7-9 ns MD simulation and used for QM/MM calculation. The experimental shifts were obtained with *E. coli* DHFR:TMP at 300 K on 800 MHz solution-state NMR spectrometer. The residues that have large difference (cs difference N > 10 ppm) are colored in red.

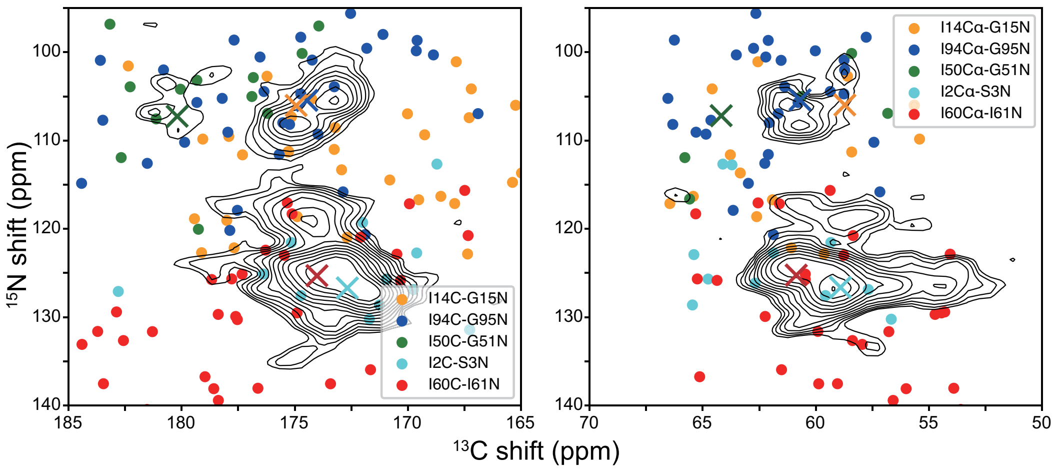


**Figure S2**: N-C correlation spectra of $^{13}$C,$^{15}$N-Ile, $^{15}$N-Gly labeled *E. coli* DHFR:TMP (IG-DHFR) were collected in DNP spectrometer and overlapped with MD/QM/MM chemical shifts predicted without local minimization before QM/MM. Restricted minimization was only applied before MD simulation on X-ray crystallography structure of DHFR:TMP (1). 100 snapshots were extracted from 1000 ns MD simulation with an interval of 10 ns and no minimization was performed before QM/MM calculation. Both C and N chemical shifts from QM/MM calculation were calibrated with the offsets from average chemical shifts of snapshots and solution-state NMR chemical shifts. The cross markers in (b) indicate the solution-state NMR chemical shifts for the 5 residue pairs.

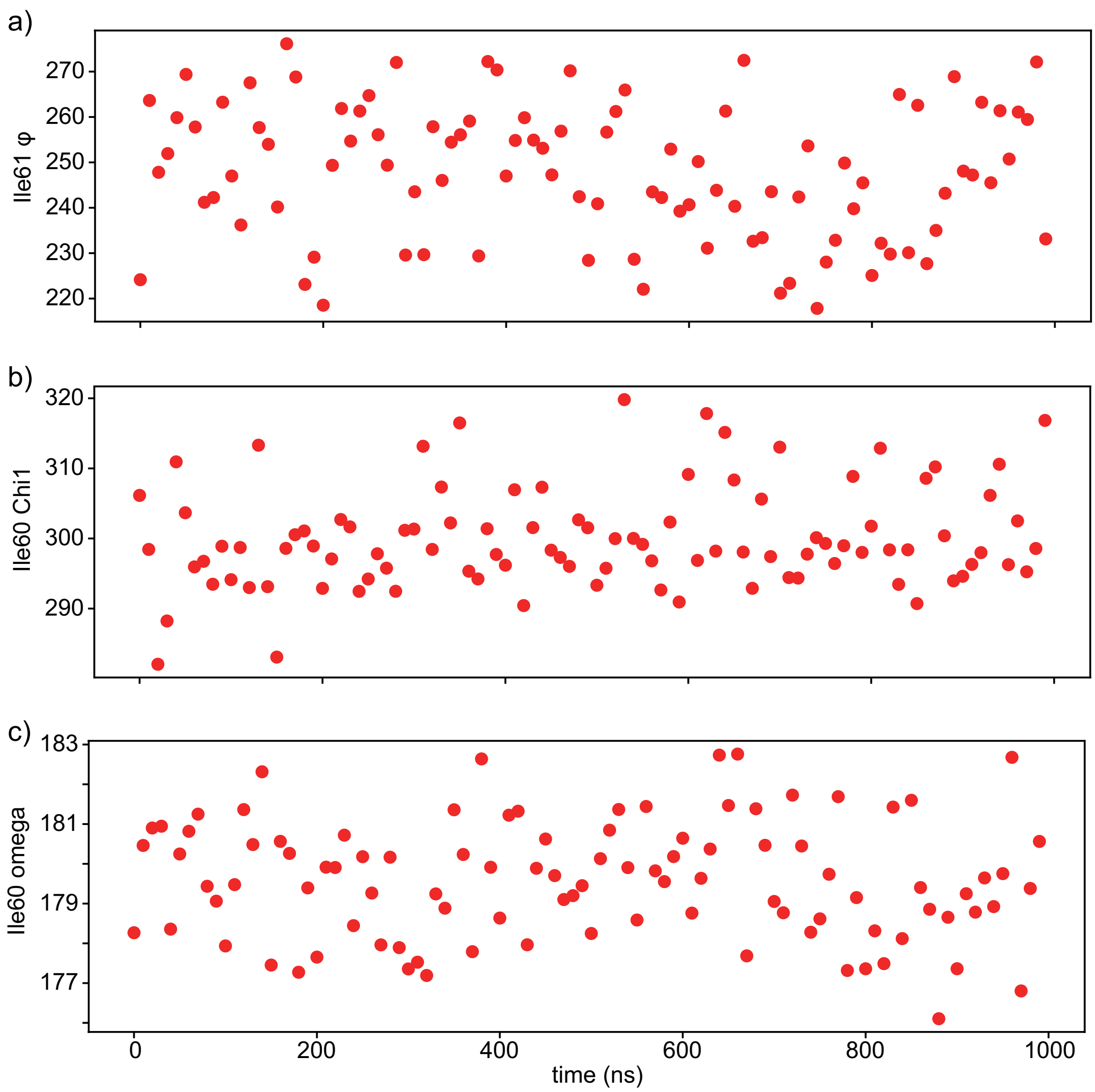


**Figure S3:** The torsion angle I61$\varphi$(a) variation during the 1000 ns MD simulation is ~ 60 degrees with snapshots extracted every 10 ns, which is similar as I60ψ. I60χ1 (b) exhibits a variation up to 40°. I60$\omega$ has a relatively smaller variation range compare to other backbone torsion angles.

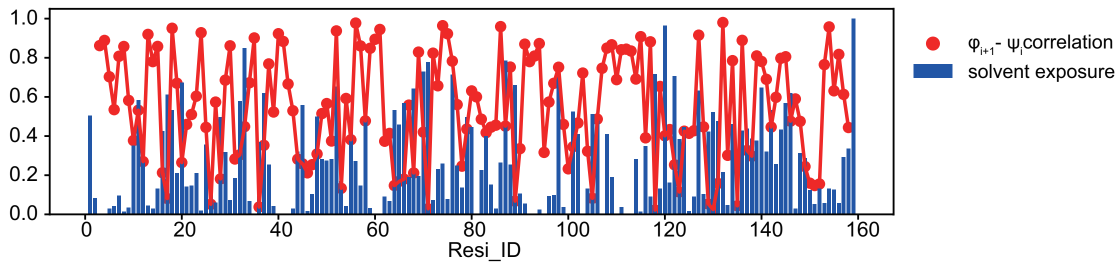


**Figure S4**: The correlation of $\varphi_{i+1}$ vs. $\psi_i$ match the pattern of solvent accessibility of each residue. The $\varphi_{i+1}$ - $\psi_i$ correlation of each residue is the correlation coefficient obtained from the 20 snapshots from 1000 ns MD simulation. The solvent exposure is normalized from the calculated solvent accessible surface area using the X-ray crystallography structure of DHFR:TMP (1), which is the same structure used for MD simulation preparation. The residues that have less solvent accessibility (normalized surface area < 0.4), like residue 2-9, 39-42, 59-61, 90-97, 108-114, 153-158, exhibit better $\varphi_{i+1}$ - $\psi_i$ correlation (r > 0.6).

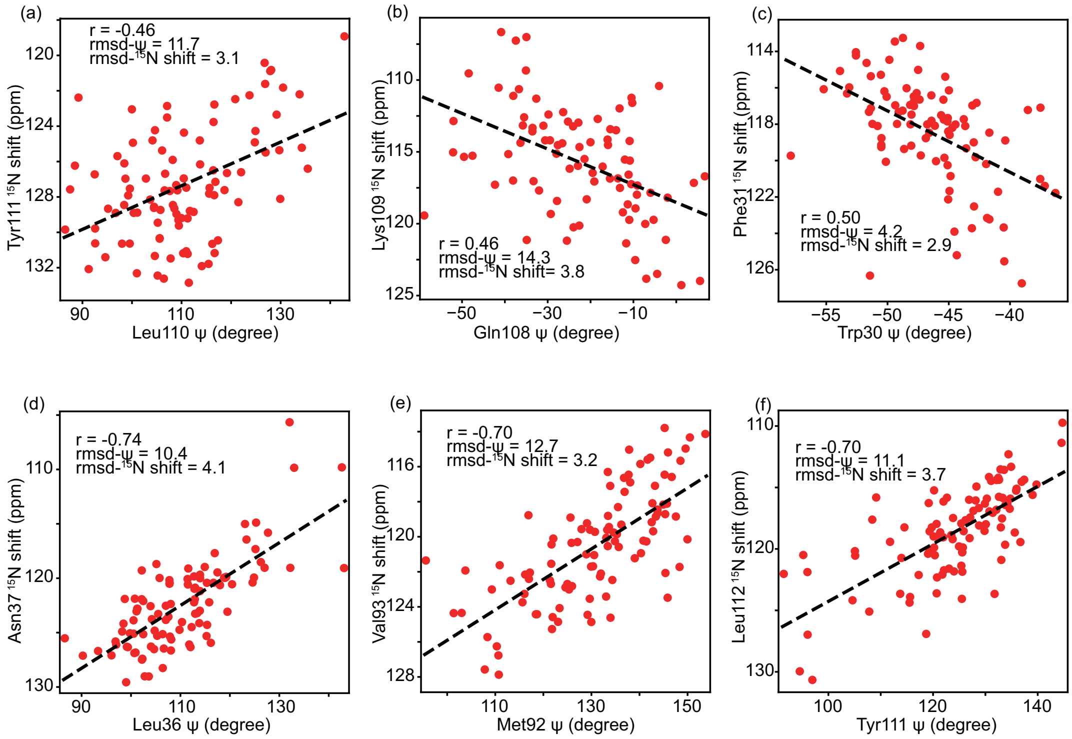


**Figure S5:** QM/MM calculated $N_{i+1}$ isotropic shifts are plotted as a function of $\psi_i$ for Tyr111-Leu110 (a), Lys109-Gln108 (b), Phe31-Trp30 (c), Asn37-Leu36 (d), Val93-Met92 (e), Leu112-Tyr111(f). 100 minimized snapshots were extracted from 1000 ns MD trajectory with an interval of 10 ns and used for QM/MM calculations. The strong correlation between $N_{i+1}$ and $\psi_i$ is not unique for Ile61N and Ile60ψ, as shown in (d), (e), (f). But clearly, not every residue exhibit this linear correlation, (a), (b), (c).

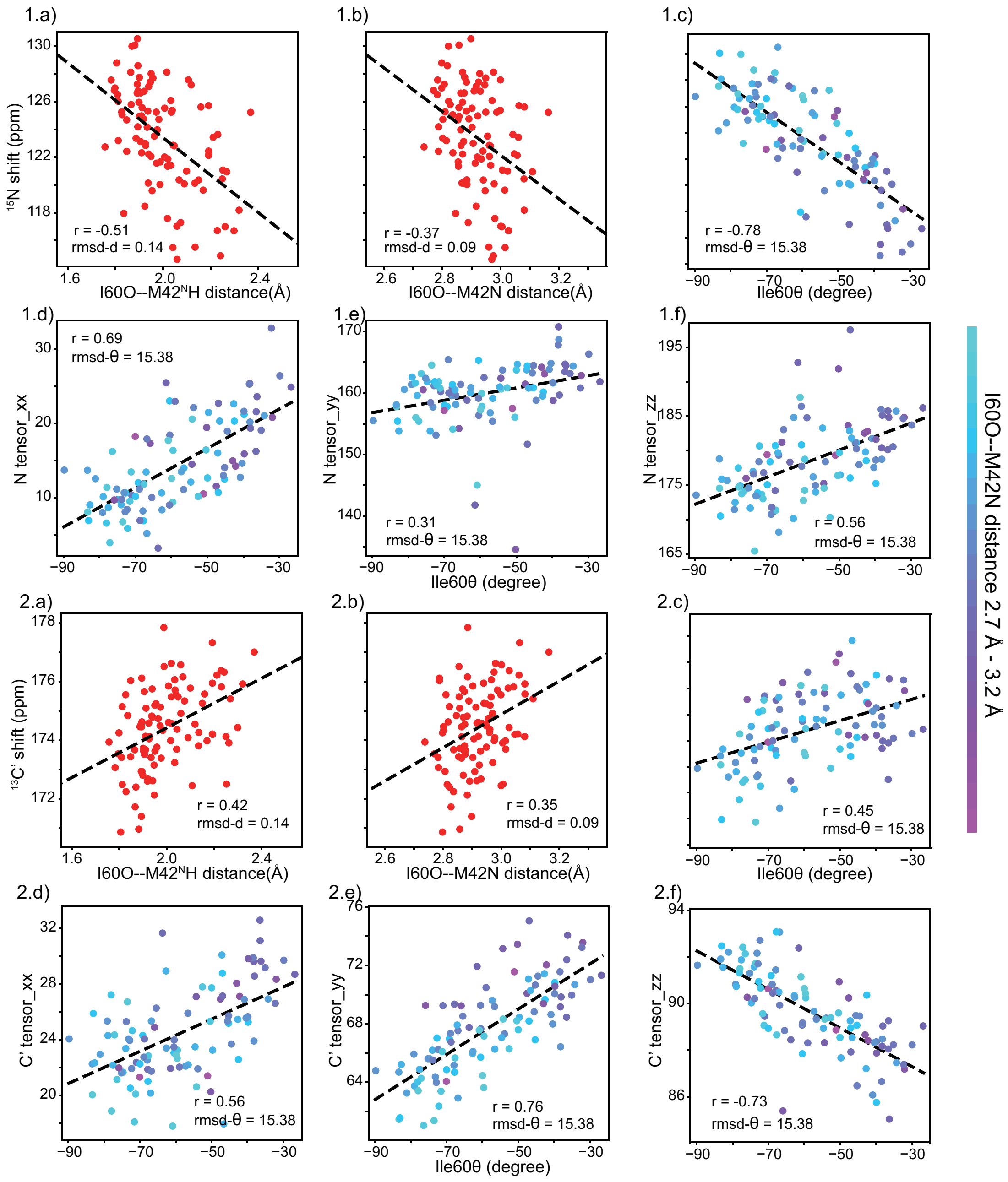


**Figure S6:** The correlation between $^{15}N$ (1.a-c) or $^{13}C'$ (2.a-c) chemical shifts and hydrogen bond distances (a) or closest heavy atom distances (b) is weaker than that with hydrogen bond angles (c). The chemical shift tensors for both $^{15}N$ (1.d-f) and $^{13}C'$ (2.d-f) show correlation with the hydrogen bond angles formed between H bond (I60O---M42$^{N}$H) and I60C = I60O.

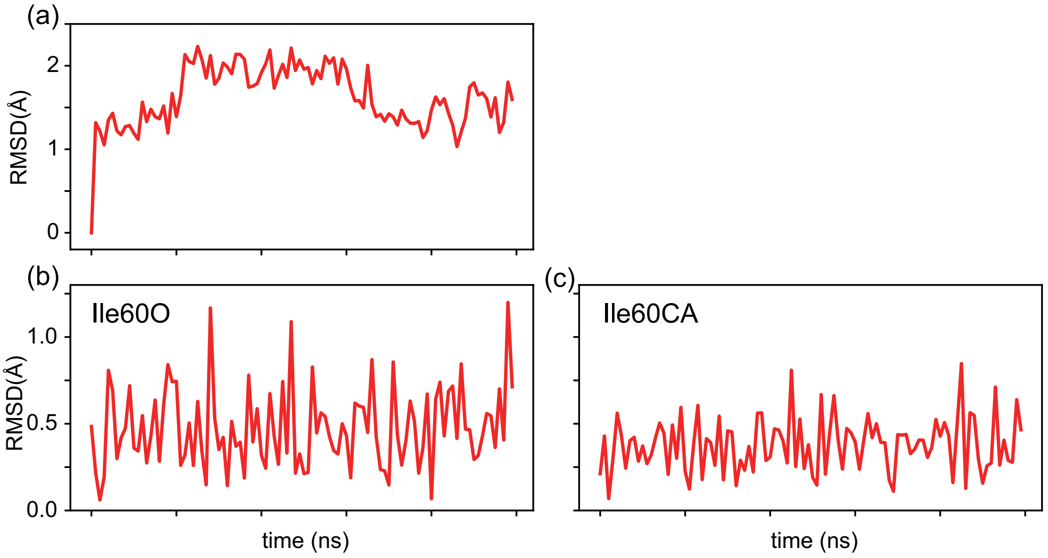


**Figure S7**: (a) The RMSD of all the backbone heavy atoms in *E. coli* DHFR referencing the initial conformation indicates that the overall protein structure preserves stability during the 1000 ns MD simulation. The RMSD of Ile60O(b), Ile60Cα(c) were calculated relative to the average structure of the 100 snapshots from 1000 ns MD simulation. Ile60Cα exhibits the least movements in the MD trajectory as it gives the smallest average RMSD ($\overline{RMSD_{I60C\alpha}}$ = 0.38, $\overline{RMSD_{I60O}}$ = 0.48). Comparing the RMSD of Ile60O(b) and Ile60Cα(c), the O undergoes a more extensive movement in the rocking motion of the amide.

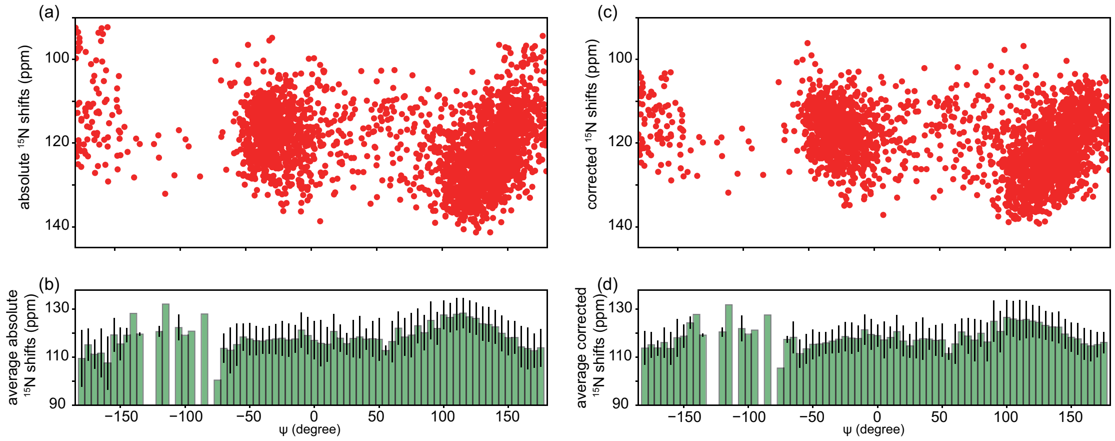


**Figure S8:** QM/MM calculated absolute (a) and corrected (c) $^{15}N_{i+1}$ chemical shifts of all the residues in 100 snapshots from 1000 ns MD simulation are plotted with their backbone torsion angle $\psi_i$. The corrected $^{15}N$ chemical shifts were referenced to the average $^{15}N$ shifts of each amino acid from BMRB (30). The corrections for each type of amino acid are listed in **Table S1**. The averaged absolute (b) and corrected (d) $^{15}N_{i+1}$ chemical shifts over 5 degrees of $\psi_i$ were shown with the standard deviations marked as error bars. The trends in the scatter plots (a, c) indicate the torsion angle has a contribution to N shifts. The well populated basin for α helices (ψ range -50°~ 0°) has a mean $^{15}N$ shift of 116 +/-5 ppm. Typical β strand residues with ψ of 100°-130° show $^{15}N$ shifts of 127 +/-6 ppm, and β strands that are closer to flat or extended conformations with ψ > 150° exhibit shifts of 115 +/-5 ppm. Notably, $^{15}N_{i+1}$ shifts vary by about 10 ppm for a narrow range of torsion angle $\psi_i$ (b, d), supporting the role of other important factors affecting the chemical shift.

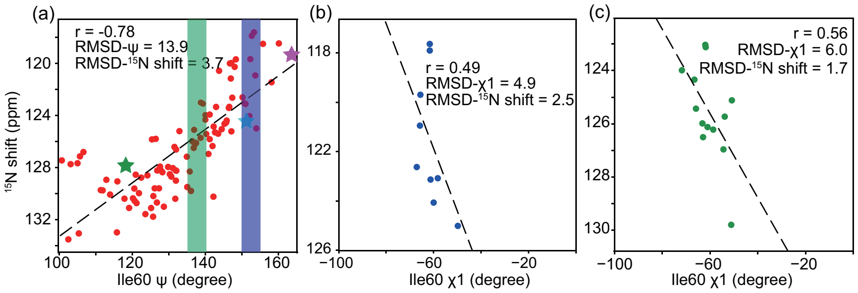


**Figure S9:** (a) QM/MM calculated Ile61N isotropic shifts are plotted as a function of I60ψ from 100 minimized snapshots from MD trajectory (red, circle). Three ψ values were measured experimentally at three $^{15}$N shift positions for Ile60 in low temperature DNP-enhanced NMR experiments (spectra shown in **Figure 2b**) and are shown here depicted with stars. The sidechain torsion angle I60χ1 were plotted with $^{15}$N shifts for snapshots with similar backbone torsion angle ψ, 150°< I60ψ <155° (b, blue), 135° < I60ψ < 140° (c, green). Ile61N shifts vary up to 8 ppm with a 5° range of I60ψ (b), and I60χ1 is likely to contribute to this variation.

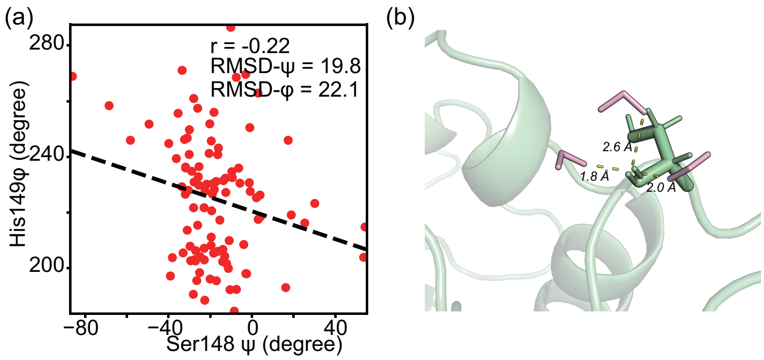


**Figure S10**: (a) The backbone torsion angles $\psi_i$ and $\varphi_{i+1}$ are more poorly correlated for residues that are exposed to solvent, for example, Ser148-His149. In the snapshot extracted at 40 ns (b) from the 1000 ns long MD simulation, three hydrogen bonds are formed between the carbonyl O of Ser148 and adjacent solvent water H. The hydrogen bond length ($O_{amide}$---$H_{water}$) ranges from 1.8 Å to 2.6 Å with various hydrogen bond angles, indicating more complicated hydrogen bond effects from solvent than protein residues.

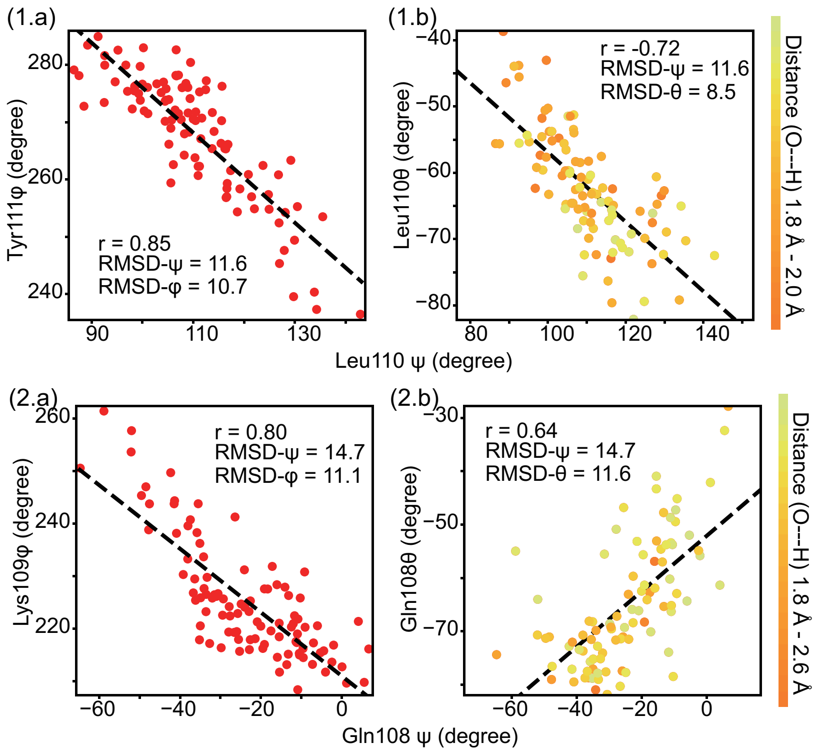


**Figure S11**: The backbone torsion angle $\psi_i$ is correlated with (a) $\varphi_{i+1}$, (b) the hydrogen bond angle $\theta_i$ at residue sites Leu110-Tyr111(1.a-b) and Gln108-Lys109 (2.a-b). All these sites form well-defined (shortest) hydrogen bonds with other protein residues and have no water molecules within 4 Å range of the amide ($O_{amide}$---$O_{water}$ > 4 Å).

**Table S1**: The $^{15}N$ chemical shift corrections were calculated based on the average chemical shifts of each kind of amino acids from BMRB (2).

| amino acid type | atom type | average chemical shift | average offset |
|---|---|---|---|
| ALA | N | 123.349 | -2.9106 |
| ARG | N | 120.816 | -0.3776 |
| ASN | N | 118.897 | 1.5414 |
| ASP | N | 120.7 | -0.2616 |
| CYS | N | 120.383 | 0.0554 |
| GLN | N | 119.943 | 0.4954 |
| GLU | N | 120.718 | -0.2796 |
| GLY | N | 109.661 | 10.7774 |
| HIS | N | 119.669 | 0.7694 |
| ILE | N | 121.443 | -1.0046 |
| LEU | N | 121.946 | -1.5076 |
| LYS | N | 121.028 | -0.5896 |
| MET | N | 120.079 | 0.3594 |
| PHE | N | 120.394 | 0.0444 |
| PRO | N | 134.565 | -14.1266 |
| SER | N | 116.29 | 4.1484 |
| THR | N | 115.383 | 5.0554 |
| TRP | N | 121.649 | -1.2106 |
| TYR | N | 120.713 | -0.2746 |
| VAL | N | 121.142 | -0.7036 |
| Ave of all aa | | 120.4384 | |
